# Single cell genotyping of matched bone marrow and peripheral blood cells in treatment naive and AZA-treated MDS and CMML

**DOI:** 10.1101/2022.01.20.476890

**Authors:** Annatina Schnegg-Kaufmann, Julie A. I. Thoms, Golam Sarower Bhuyan, Henry R. Hampton, Lachlin Vaughan, Kayleigh Rutherford, Purvi M. Kakadia, Emma M. V. Johansson, Tim Failes, Greg M. Arndt, Jason Koval, Robert Lindeman, Pauline Warburton, Alba Rodriguez-Meira, Adam J. Mead, Ashwin Unnikrishnan, Stefan K. Bohlander, Elli Papaemmanuil, Omid Faridani, Christopher J. Jolly, Fabio Zanini, John E. Pimanda

## Abstract

Progressively acquired somatic mutations in hematopoietic stem cells are central to pathogenesis in myelodysplastic syndromes (MDS) and chronic myelomonocytic leukemia (CMML). They can lead to proliferative advantages, impaired differentiation and progressive cytopenias. MDS or CMML patients with high-risk disease are treated with hypomethylating agents including 5-azacytidine (AZA). Clinical improvement does not require eradication of mutated cells and may be related to improved differentiation capacity of mutated hematopoietic stem and progenitor cells (HSPCs). However, the contribution of mutated HSPCs to steadystate hematopoiesis in MDS and CMML is unclear. To address this, we characterised the somatic mutations of individual stem, progenitor (common myeloid progenitor, granulocyte monocyte progenitor, megakaryocyte erythroid progenitor), and matched circulating (monocyte, neutrophil, naïve B cell) haematopoietic cells in treatment naïve and AZA-treated MDS and CMML via high-throughput single cell genotyping. The mutational burden was similar across multiple hematopoietic cell types, and even the most mutated stem and progenitor clones maintained their capacity to differentiate to mature myeloid and, in some cases, lymphoid cell types *in vivo.* Our data show that even highly mutated HSPCs contribute significantly to circulating blood cells in MDS and CMML, prior to and following AZA treatment.

**Key points:**

* Highly mutated HSPCs contribute significantly to circulating blood cells in MDS and CMML, prior to and following AZA treatment.
* The mutational burden in matched bone marrow and peripheral blood cells in MDS and CMML was similar throughout myelopoiesis.

## INTRODUCTION

Somatic mutations in hematopoietic stem cells (HSCs) are a central pathogenic event in myelodysplastic syndromes (MDS) and chronic myelomonocytic leukemia (CMML) where they induce proliferative advantages and impaired differentiation with subsequent cytopenias in peripheral blood (PB)^1–6^. Patients with high-risk disease who are ineligible for transplantation are treated with hypomethylating agents, usually 5-azacytidine (AZA). AZA treatment can improve PB counts and delay progression to AML in some patients^7–9^. However, AZA response does not require eradication of mutated HSCs. We and others have previously described cohorts with haematological response to AZA despite persistently high variant allele fractions in bone marrow (BM)^10,11^. Colonies derived from *in vitro* assays of stem cell function following AZA treatment showed decreased mutational complexity, suggesting a shift in haematopoiesis from clones with high to low mutational burden in response to treatment^11^. However, *in vitro* colony-forming capacity might not correlate with *in vivo* hematopoietic potential, and the relative contribution of high vs low/unmutated HSCs to circulating blood cell types *in vivo* remains unknown. Single cell genotyping techniques can be used to resolve combinations of mutations in *ex vivo* cells^1,12–15^ However, to our knowledge such techniques have not been applied to resolve the relative contributions of stem and progenitor cells with multiple mutations to circulating mature blood cells in MDS/CMML. Here we use a combination of index sorting and single cell genotyping to characterise the haplotype composition of individual stem (HSC/multipotent progenitors (MPP), MDS stem cells (MDS-SC), progenitor (common myeloid progenitor (CMP), granulocyte monocyte progenitor (GMP), megakaryocyte erythroid progenitor (MEP)), and high-turnover circulating cells (monocyte, neutrophil, naïve B cell (nBC)) in treatment naïve and AZA-treated MDS and CMML.

## METHODS

BM and PB samples were collected with patient consent and institutional ethics approval. BM was enriched for CD34+ cells and target populations single-cell index sorted into 384-well plates (Figure S1). Target mutations were amplified in single cells by modifying TARGET-Seq^15^ followed by barcoding and Illumina sequencing (Figure S2; Supplemental Material). Capture sequencing was performed using a targeted panel for myeloid driver mutations (Supplementary Material). Raw data is available at SRA (accession: PRJNA798507) and https://flowrepository.org/id/FR-FCM-Z4PR. Mutational calling was performed via pairwise sequence alignments using SeqAn^16^ and seqanpy (https://github.com/iosonofabio/seqanpy). Analysis code is available at https://github.com/julie-thoms/MDS_amplicons.

## RESULTS and DISCUSSION

We analysed matched stem and progenitor cells from BM and high-turnover differentiated cells from PB in three patients (Figure 1A, B, Table S1). Variant allele fractions (VAFs) were determined by capture sequencing in bulk samples from each cell type. The distribution of VAFs were similar across BM and PB cell types in all patients with the notable exception that nBCs in patient H198304 were predominantly wild-type (wt) (Figure 1C). We confirmed this exception using an orthogonal approach (Figure S3). We then modified TARGET-Seq to determine VAFs of these known mutations in single cells (Figures 1D, S4). Allele fractions were highly correlated between bulk- and single-cell assessments (Figure S5; Pearson r = 0.8989).

**Figure 1.**
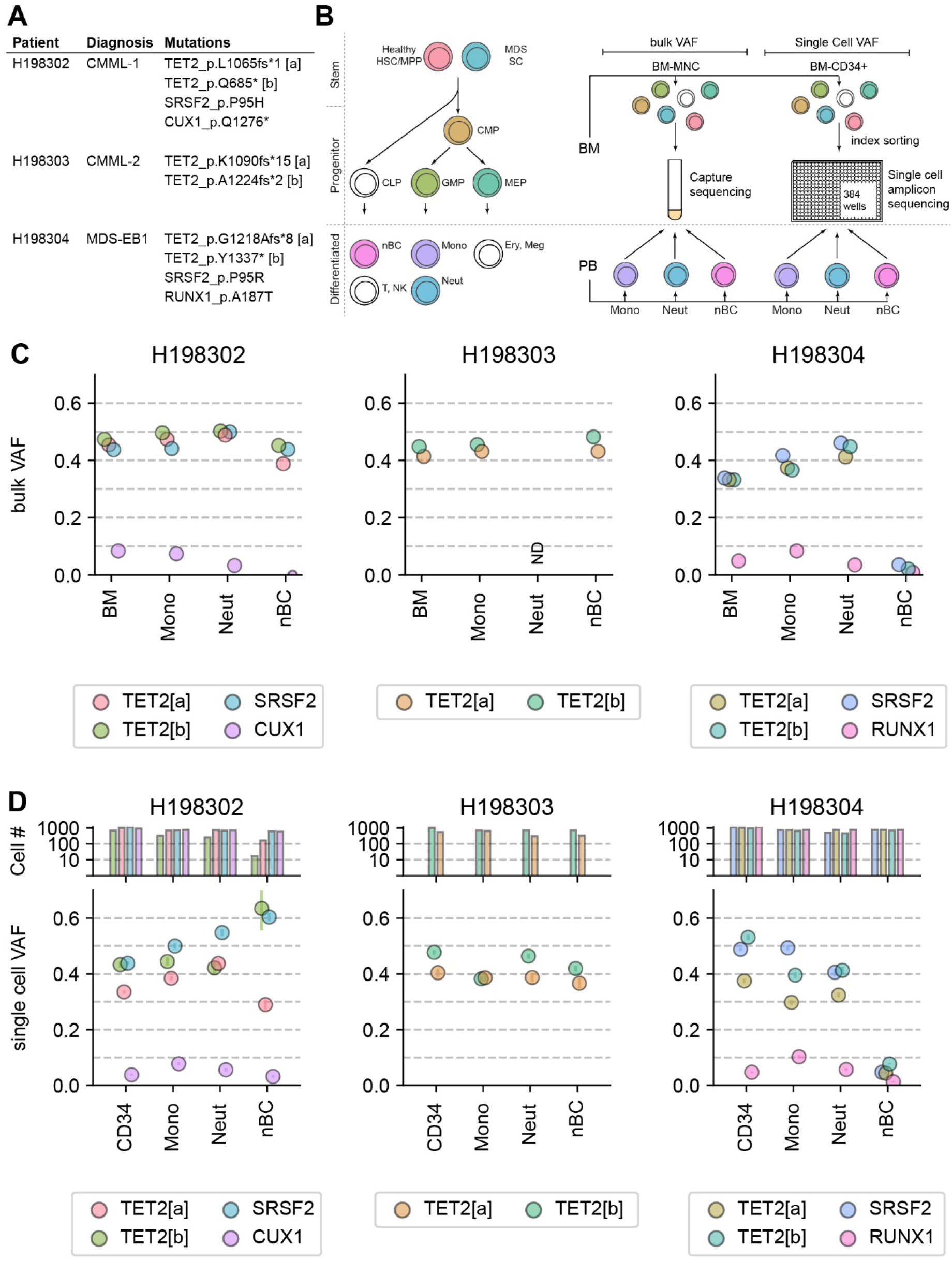
Mutational burden in bone marrow progenitors and differentiated peripheral blood cells. **(A)** Diagnosis and tracked mutations in the three patients included in this study. **(B)** *Left:* Blood differentiation hierarchy in MDS/CMML showing stem (healthy stem cells [HSC/MPP] and MDS stem cells [MDS-SC]), progenitor (CMP, GMP, MEP, CLP), and differentiated mature cells. Cell types coloured white were not characterised in this study. *Right:* Schematic showing collection and cell sorting strategies for peripheral blood (PB) and bone marrow (BM). PB was flow sorted into neutrophils (Neut: SSC^lo^, CD45^+^, IgD^-^, CD16^+^, CD66b^+^), monocytes (Mono: SSC^lo^, CD45^+^, IgD^-^, CD16^+^), and naïve B cells (nBC: SSC^lt5^, CD45^+^, IgD^+^, CD27^-^). Bone marrow mononuclear cells (BM-MNCs) were isolated on Ficoll and used directly for bulk capture sequencing. MACS-enriched CD34+ cells (BM-CD34+) were dropped into 384 wells plates for amplicon sequencing, with indexing for CD38, CD 123, CD45RA, CD90, and IL1RAP and post-hoc assignment of cell type (HSC/MPP: LIN^-^, CD34^+^, CD38^lo^, CD45RA^-^, CD123^-^, IL1RAP^-^; MDS-SC: LIN^-^, CD34^+^, CD38^lo^, [CD45RA^+^ or CD123^+^ or IL1RAP^+^]; CMP: LIN^-^, CD34^+^, CD38^+^, CD45RA^-^, CD123^+^; GMP: LIN^-^, CD34^+^, CD38^+^, CD45RA^+^, CD123^+^; MEP: LIN^-^, CD34^+^, CD38^+^, CD45RA^-^, CD123^-^) **(C)** Variant allele fraction (VAFs) determined by capture sequencing in bulk BM and PB cell types. **(D)** VAFs determined by amplicon sequencing in single cells. Each allele was analysed individually. Upper bar graph indicates the number of cells analysed for each mutation in each cell type. Lower graph shows average VAF (i.e., the proportion of reads containing the known mutation) across all cells with at least 10 reads mapping to that amplicon. Bars show standard error of the mean (sem).

Stem cells with multiple mutations (Figure 2A) could be subject to negative clonal selection with a bias towards less mutated stem and progenitor cells contributing to circulating differentiated cells, or neutral selection with similar contributions irrespective of mutational burden (Figure 2B). To resolve this, we first classified BM cells from each of the three patients into healthy HSC/MPP, MDS-SC, CMP, GMP, or MEP using indexed fluorescence activated cell sorting (FACS) and assessed the presence/absence of known variants in each of these BM cell types and in matched PB neutrophils, monocytes and nBCs (Figure 2C-E).

**Figure 2.**
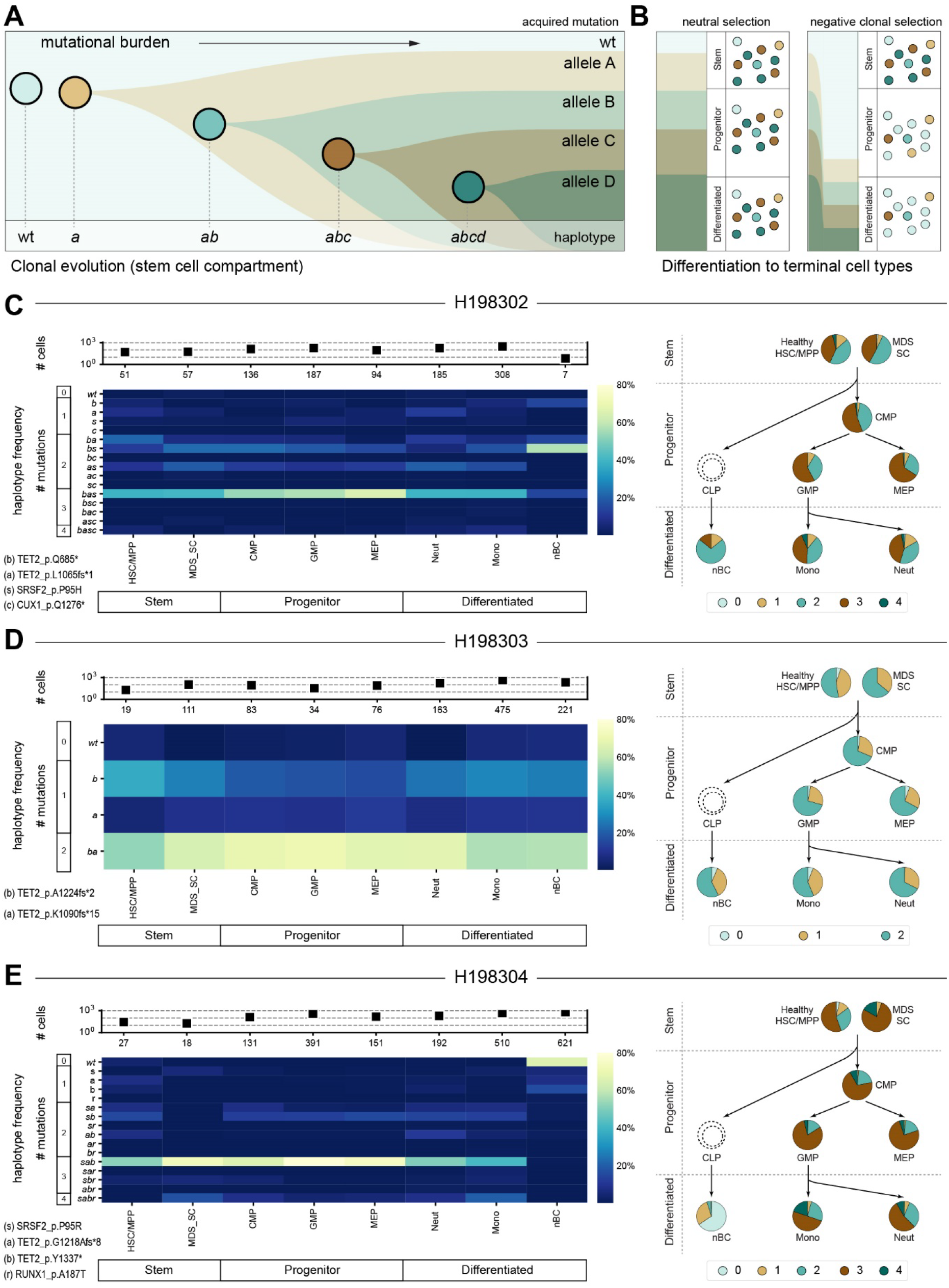
Fitness of highly mutated MDS and CMML stem cells to produce terminal blood types. **(A)** Schematic showing hypothetical clonal evolution in MDS/CMML with sequential acquisition of mutations in four alleles (allele A, allele B, allele C, allele D; note that in some cases both alleles of a single gene may independently acquire mutations e.g., in the case of *TET2*). The combined genotype (haplotype) of each resulting cell population is indicated, with lower case letters indicating the presence of a mutated allele. **(B)** Models of terminal blood production in MDS/CMML. In the neutral selection model *(left)* stem cells with multiple mutations retain capacity to produce terminal blood types, although there may be a reduction in absolute cell number or functionality. In the negative clonal selection model (*right*), cells harbouring multiple mutant alleles are abundant in the stem cell compartment but have reduced differentiation capacity resulting in a higher frequency of wild-type cells in the circulating population. **(C-E)** Single cell haplotypes in three patients. Heatmaps *(left panels)* show the observed frequency of all combinations of mutations, while pie charts *(right panels)* show the proportions of cells across the haematopoietic hierarchy carrying zero, one, two, three, or four mutations in the specified alleles. **(C)** Patient H198302. Alleles shown are TET2_p.Q686* (*b*), TET2_p.L1065fs*1 *(a),* SRSF2_p.P95H (*s*), CUX1_p.Q1276* (*c*). **(D)** Patient H198303. Alleles shown are TET2_p.A1224fs*2 (*b*), TET2_p.K1090fs*15 *(a).* **(E)** Patient H198304. Alleles shown are SRSF2_p.P95R (*s*), TET2_p.G1218fs*8 *(a),* TET2_p.Y1337*, (*b*) RUNX1_p.A187T (*r*).

In patient H198302 (CMML; AZA therapy ~ 10 years; complete responder (IWG^17^); Table S1), we tracked four mutations (Figure 2C; *SRSF2, CUX1* and biallelic *TET2)* with acquisition order inferred from VAF at diagnosis^11^. Most stem cells carried two or three of the tracked mutations and we did not detect any wt HSC/MPP. Cells across the progenitor compartment were similar and mostly highly mutated – a pattern that was maintained particularly in differentiated monocytes and neutrophils with a substantial proportion of the circulating cells derived from highly mutated progenitors.

In patient H198303 (CMML; AZA therapy ~ 10 years; complete responder (IWG^17^); Table S1), we tracked two mutations detected at diagnosis^11^ (Figure 2D; biallelic *TET2).* In the stem compartment, cells carrying biallelic *TET2* mutations were frequent, but numerous cells carried either a single mutant allele or were wt. Progenitors were relatively homogenous, although mutational burden differed slightly from the stem compartment. The haplotype distribution in differentiated cells was again similar to progenitor cells, with the exception that we did not detect any wt neutrophils.

In patient H198304 (MDS-EB1; no HMA therapy), we tracked four mutations (Figure 2E; *SRSF2, RUNX1* and biallelic *TET2).* Most stem cells carried two or three mutations, a few healthy HSC/MPPs had no mutations detected, and around 20% of MDS-SC carried an additional mutated *RUNX1* allele. In the progenitor compartment all cell types were similar; most cells carried three mutations. In the differentiated compartment, myeloid populations contained cells with one, two, three, or four mutations, with a higher proportion of cells with all four alleles mutated than progenitors, suggesting that highly mutated stem cells can produce differentiated myeloid cells, and that cells carrying the additional *RUNX1* mutation are advantaged in this regard compared to cells with *TET2* and *SRSF2* mutations alone. RUNX1_p.A187T lies within the DNA- and protein-binding runt homology domain and may alter transcriptional activity. Conditional *RUNX1* knockout leads to myeloid proliferation^18^; the patient’s mutation may similarly favour myeloid expansion or differentiation. Consistent with bulk analysis, very few mutant cells were detected in nBCs, suggesting that the small wt

HSC/MPP population is the predominant origin of nBCs in this individual. Analysis of additional lymphoid populations in PB from this patient revealed that naïve T, but not NK, cells were also predominantly wt (Figure S3C-D), suggesting specific impairment of B- and T-lineage maturation in the mutated cells.

All patients had clones with biallelic *TET2* mutations, with or without additional oncogenic mutations, and shared the following features: (a) Very few, if any, sampled HSCs (with or without aberrant CD45RA^+^ or CD123^+^ or IL1RAP^+^) were wild-type, (b) A high proportion of sampled HSCs and myeloid progenitors in the bone marrow had two or more mutations, (c) Few, if any of the sampled circulating neutrophils or monocytes were wild-type, (d) The mutation profiles of PB neutrophils and monocytes (very high-turnover cells) mirrored those that were present in corresponding myeloid progenitors in the bone marrow. Attrition of highly mutated cells during myeloid maturation was not observed in any of the three patients, irrespective of HMA therapy, suggesting that *in vivo*, highly mutated stem and progenitor cells retain the capacity to differentiate. However, there were individual differences. In particular, in patient H198304 the rarity of mutated nBC compared to mutant NK and myeloid cells might be attributed to the order of mutation acquisition^19–22^, the presence of an additional undetected somatic mutation in mutant cells that specifically impacts B cells, or individual differences in the bone marrow microenvironment^23,24^.

In summary, we characterised the mutational profile of thousands of individual stem, progenitor, and differentiated cells from three MDS/CMML patients and found that *in vivo,* highly mutated stem cells contribute to the circulating population prior to and following AZA treatment and were preferentially poised towards specific differentiation trajectories. These findings are particularly relevant in an era of combining cytotoxic therapies designed to eliminate mutant cells with hypomethylating agents^25^. Our findings emphasise that therapeutic principles and endpoints that apply in high-blast AML (clonal eradication and minimal residual disease monitoring for relapse) may not be appropriate when treating high-risk MDS/CMML where clinical response does not require eradication of mutant clones; indeed, eradication may be detrimental. Further studies will help determine the prevalence of this phenomenon among cohorts of MDS and CMML patients with different mutation profiles and whether it correlates with clinical outcomes.

## ACKNOWLEDGEMENTS

We thank study participants for their generosity in providing samples, and Swapna Joshi and Ameline Lim for assistance in processing PB and BM. AS-K was supported by a Swiss national research foundation early postdoc mobility grant (P2LAP3_181273). SKB and PMK are supported by Leukaemia and Blood Cancer New Zealand and the family of Marijana Kumerich. JEP is supported by project grants from the National Health and Medical Research Council of Australia (APP1042934, APP1102589, APP1008515), a translational program grant from the Leukemia Lymphoma Society (LLS)-Snowdome Foundation-Leukaemia Foundation, project funds from the Translational Cancer Research Network – a Translational Cancer Research Centre funded by the Cancer Institute NSW, Anthony Rothe Memorial Trust, and philanthropic funding from Christina’s Light.

## AUTHORSHIP CONTRIBUTIONS

AS-K, JAIT, RL, PW, AR-M, AJM, AU, OF, CJJ, FZ, JEP designed research.

AS-K, JAIT, GSB, HRH, LV, PMK, EMVJ, TF, GMA, JK, OF, CJJ performed research.

AS-K, JAIT, KR, PMK, SKB, EP, CJJ, FZ, JEP analyzed and interpreted data.

AS-K, JAIT, CJJ, FZ, JEP wrote the manuscript.

