## Supplementary methods and figures for "Single cell genotyping of matched bone marrow and peripheral blood cells in treatment naive and AZA-treated MDS and CMML"

Details of commercial reagents are in Table S2.

##### Patients

This study investigated three patients (Figure 1A, Table S1). Two of these patients (H198302, H198303) were subjects in a previous clinical trial[1] and have both been treated long term with azacytidine (AZA) (Table S1). A third patient (H198304) was relatively recently diagnosed with MDS-EB1 and has never been treated with AZA. Peripheral blood and bone marrow samples were collected with patient consent in accordance with the Declaration of Helsinki, and with institutional ethics approval ref:08/190 from South Eastern Sydney Local Health District, NSW, Australia.

##### Preparation of bone marrow (BM) and peripheral blood (PB) mononuclear cells (MNCs)

BM (30mL) was collected on 10mL Hanks balanced salt solution (Invitrogen) containing Heparin (500U) and stored at 4ºC with rocking for up to 48 hours after collection. BM was diluted 1:5 in RPMI (Invitrogen) and MNCs enriched via density gradient separation with Lymphoprep (Eli Tech) at 800g/room temperature/30 min with low brake and resuspended in AutoMACS running buffer (Miltenyi Biotec). Aliquots of $10^6$ cells were spun down and the pellets frozen and stored at -80ºC for later DNA extraction and bulk variant allele fraction (VAF) analysis. Remaining BM MNCs were enriched for CD34+ cells using MACS CD34 microbeads and an AutoMACS separator (both Miltenyi Biotec) according to manufacturer instructions and resuspended in PBS at $10^7$/mL.

PB was collected in EDTA tubes and stored at room temperature on a shaker for up to 24 hours. Whole blood was centrifuged at 800g/room temperature/10 min, and the cell pellet resuspended in 15ml of red cell lysis buffer (Pharmingen) and incubated at room temperature with rocking for 15 min. After two rounds of red cell lysis cells were resuspended in PBS at $10^7$/mL.

**Cell staining**

Isolated BM and PB cells were stained 4°C/20 min with Zombie Yellow (Biolegend) diluted 1/500, then staining quenched by the addition of 1-2 volumes of FACS buffer (PBS containing 5% BSA and 2mM EDTA). Cells were then centrifuged and stained with the either the BM staining panel (Table S3) or the PB staining panel (Table S4) at 4°C/20 min, diluted in 2-3 volumes of FACS buffer, and filtered prior to sorting. A supplementary sort targeting lymphocytes and monocytes was performed with blood collected the previous day from patient H198304, using DAPI exclusion in place of Zombie Yellow to identify viable cells (Table S4).

**Cell sorting**

Single cells were index-sorted with a FACSAria™ III (Becton Dickinson) into 384-well plates containing 2µL of lysis buffer (0.025% TritonX-100 and 0.60 units/µL recombinant RNase inhibitor). Representative gating is shown in Figures S1A and S1B. From BM we sorted $LIN^-$, $CD34^+$ and a smaller number of $LIN^-$, $CD34^+$, $CD38^-$ cells to increase the representation of early stem cells which are relatively rare in the $CD34^+$ pool. From PB we sorted neutrophils (Neut: $SSC^{hi}$, $CD45^+$, $IgD^-$, $CD16^+$, $CD66b^+$), monocytes (Mono: $SSC^{lo}$, $CD45^+$, $IgD^-$, $CD16^+$), and naïve-enriched B cells (nBC: $SSC^{lo}$, $CD45^+$, $IgD^+$, $CD27^-$). Plates of sorted cells were sealed, centrifuged at 2000g/4°C/30 sec, and placed on dry ice prior to storage in vapour phase of liquid nitrogen. FCS files for indexing are listed in Table S5 and are available from https://flowrepository.org/id/FR-FCM-Z4PR. PB cells (up to 5 x $10^5$ for each cell type) were bulk sorted using the same gating strategy, pelleted, and stored at -80°C for later DNA extraction and bulk VAF analysis.

In the supplementary sort of patient H198304 PB (Figure S3A), B cells were gated as small singlet DAPI$^-$ CD3$^-$ CD56$^-$ CD235ab$^-$ CD14$^-$ CD19$^+$ cells, then sorted into a IgD$^+$ CD27$^-$ (naïve-enriched) bulk population (3.8x$10^4$ cells) or into a CD27$^+$ IgD$^-$ (memory) bulk population (2.1x$10^4$ cells). A monocyte population (7.1x$10^5$ cells) was sorted with the same staining panel as large singlet DAPI$^-$ CD3$^-$ CD56$^-$ CD235ab$^-$ CD19$^-$ CD33$^+$ CD14$^+$ cells. Using a separate stain panel, NK cells were sorted as small singlet DAPI$^-$ CD19$^-$ CD235ab$^-$ CD3$^-$ CD56$^+$ cells (1.9x$10^5$ cells), naïve T cells sorted as small singlet DAPI$^-$ CD19$^-$ CD235ab$^-$ CD56$^-$ CD3$^+$ CCR7$^{hi}$ CD45RA$^{hi}$ cells (2.0x$10^5$ cells) and memory T cells sorted as small singlet DAPI$^-$ CD19$^-$ CD235ab$^-$ CD56$^-$ CD3$^+$ CD45RA$^-$ cells (2.6x$10^5$ cells). Cells were pelleted at 500*g*, washed with cold PBS twice, vortexed into suspension in residual cold PBS, then frozen on dry ice and stored overnight at -80 °C. Cell pellets were thawed with pipetting into 0.1 mL of 0.25 mg/mL proteinase k (Roche) in 1% Tween 20, 10 mM TrisHCl pH 8.0 and 0.1 mM EDTA, incubated at 56°C for 40 min to digest proteins, heated at 95°C for 10 min to inactivate proteinase k, then stored at -80°C for future use.

**Bulk VAF Analysis**

DNA was extracted from frozen pellets using either an AllPrep DNA/RNA Mini Kit or Micro Kit (Qiagen) according to manufacturer's instructions, and DNA yield and quality assessed using a Nanodrop spectrophotometer (ThermoFisher) and a Quant-iT™ PicoGreen™ dsDNA Assay Kit (Invitrogen). Capture sequencing library construction and sequencing was performed at Memorial Sloan Kettering Cancer Center and the University of Auckland using standard clinical myeloid capture arrays and reporting. The genes covered in the capture panel are in Table S6.

#### Supplementary lymphocyte VAF analysis

DNAs[2] (5 µL) from proteinase k-digested lymphocyte or monocyte populations supplementary bulk-sorted from patient H198304 PB were used as templates in 50 µL PCR reactions containing 1x Platinum *Taq* PCR buffer (Invitrogen), 2.5 mM $MgCl_2$, 0.2 mM dNTPs, 100 µM primary multiplex PCR primers (Table S7 see "Targeted primer design" below) and 2 units of Platinum *Taq* DNA polymerase. Reactions were heated at 95°C for 4 min, then cycled 10 times at 95°C for 15 s, 65–55°C with a reduction of 1°C per cycle for 30 s, then 68°C for 1 min. This was immediately followed by 25 cycles of 95°C for 15 s, 55°C for 30 s, then 68°C for 1 min. Amplicons were quantified using a Fragment Analyzer 5200 (Agilent), depleted of enzyme, dNTPs and primers using a Wizard SV Gel and PCR product Cleanup System (Promega), then sequenced using a single nested primer specific for each gene of interest to prime multiple separate Sanger sequencing reactions (Macrogen, S Korea) to determine the presence or non-detection of H198304-specific variant alleles.

#### Quantitation of antigen-induced somatic *Ig* hypermutation in B cells from patient H198304 PB

It was conceivable that non-mutated $IgD^+$ $CD27^-$ B cells in patient H198304 represented a non-canonical memory population or an expanded clone. To investigate further, we sorted naïve versus memory B and T cells, NK cells and monocytes from a fresh PB sample, and in B cells sequenced for the *IGH* somatic mutations that typify memory B cells (Fig S3). VDJ-rearrangements present in "naïve" $CD19^+$ $IgD^+$ $CD27^-$ B cells or "memory" $CD19^+$ $IgD^-$ $CD27^+$

B cells supplementary bulk-sorted from patient H198304 PB were amplified in 25 µL reactions using 5 µL input template, 500 µM FR3 consensus forward primer (JP1833; CACGGCYGTGTATTACTGTGC; Table S7) and a reverse primer upstream from JH5 (JP1835; AGGACCCAGGCAAGAAC; Table S7) [3] at 98ºC for 10s, 65ºC for 30s and 72ºC for 30s for 35 cycles using 1x Q5 HiFi PCR Mastermix (NEB). The products were purified using a Wizard SV Gel and PCR Product Cleanup System (Promega), A-tailed using 1 unit Platinum *Taq* DNA polymerase in 1x Platinum buffer (Invitrogen), 1.5 mM $MgCl_2$, 0.2 mM dATP at 72ºC for 30 min, then re-purified. Products were ligated into pGEM-T Easy vector (Promega) and transformed into competent α-select *E. coli* (Bioline) according to manufacturer instructions. The plasmid inserts from random ampicillin-resistant colonies were re-amplified by touching random single colonies into 96-well 25µL PCR reactions containing 1 unit Platinum *Taq* DNA polymerase in 1x Platinum reaction buffer (Invitrogen), 1.5 mM $MgCl_2$, 0.2 mM dNTPs, 100µM JP1833 primer and 100µM JP1835 primer, using 10 touchdown cycles of 95ºC for 15s, 65–55ºC for 30s (dropping 1ºC per cycle) and 68ºC for 2 min, followed by 25 cycles of 95ºC for 15s, 55ºC for 30s and 68ºC for 2 min. Amplicon sizes and yields were determined using a Fragment Analyzer 5200 (Agilent). Primers and dNTPs were degraded using exonuclease 1 and shrimp alkaline phosphatase [4] (NEB) and amplicons were sequenced using primer JP1835 in 96-well fluorescent Sanger sequencing reactions (Macrogen, S Korea). Sequencing traces were aligned to Homo sapiens chromosome 14 GRCh38 region 105853215-105875260 and arranged into consensus trees (to identify and exclude duplicate VDJ-recombinations) using Geneious Prime software (64 bit Java Version 11.0.11+9, Biomatters Ltd). A total of 54 quality unique sequences trimmed of coding regions (potentially mutated prior to antigen activation by error-prone non-homologous end-joining during VDJ-recombination), of insertions, and of a single SNV that differed between patient H198304's two germline *IGH* alleles were uploaded to http://shmtool.montefiore.org [5] as multiple sequence FASTA files to quantify point mutations. Deletions and insertions were collated manually from the Geneious Prime alignments. Data plotting and a two-tailed Mann-Whitney test used Prism v9.2.0 for macOS (Graphpad Software, LLC).

**Targeted primer design**

Somatic mutations in patients H198302 and H198303 were identified in a previous study[1], and mutations in patient H198304 were identified in a routine clinical myeloid mutation panel. Mutations known or suspected to be drivers of myeloid malignancy were selected for single cell genotyping (Table S1, Figure 1A). Primers were designed to amplify gDNA and cDNA across known mutation sites using a nested strategy to amplify and barcode multiple alleles in each cell (Figure S2). Targeted primers contained a phosphorothioate modification just prior to the 3' end to prevent primer degradation by the 3'-5' exonuclease activity of the DNA polymerase. In some cases, primer sets were able to distinguish between gDNA and cDNA, while other primer sets amplified both cDNA and gDNA. Primers for the primary amplification reaction were first tested in single-plex reactions using DNA from healthy cells as a template. Primer sets yielding the appropriate amplicons were then tested in multiplex assays, and those sets with suitable multiplexed behaviour were further tested in pseudo-single cell PCR reactions (50pg/reaction). Only primer sets that passed all rounds of testing were used for single cell genotyping. Nesting primers underwent similar testing with PCR products from the primary PCR used as the test template.

#### Library preparation

##### *Liquid handling*

Reagents for protease digestion, reverse transcription and preamplification were added to each well using a Mantis (r) Liquid Handler (Formulatrix) or a mosquito LV (SPT Labtech). PCR reactions were transferred between plates using a Janus liquid handling robot (Perkin Elmer) or a mosquito LV (SPT Labtech). Barcoding master plates where each well contained one unique combination of barcoding primers were prepared using an EP-Motion (Eppendorf) liquid handling robot system.

##### *Protease digestion and reverse transcription*

Plates of single sorted cells were thawed by centrifugation at max speed/4°C/2 min, and 1µL of protease reagent (25 mM Tris-Cl pH8.3, 0.05AU/mL protease (Qiagen), 5 µM custom Oligo dT (JP1745; Table S7), 2.5mM/each dNTP (Bioline)) added to each well. Plates were briefly mixed and centrifuged, then incubated at 37°C/10min, 70°C/10min then placed on ice. 2 µL of RT-reagents (112.5mM Tris pH 8.3, 175mM NaCl, 1.25mM GTP, 6.25mM MgCl, 12.5 mM DTT, 1U/µL RNAse Inhibitor, 5µM TSO-LNA (JP1692; Table S7), 12.5U/µL Maxima H Minus reverse transcriptase) were added to each well, plates spun down, mixed, and spun down again, and then incubated at 42°C/90min, 10 cycles of 42°C/2min and 50°C/2min, then 85°C/5min before being placed on ice.

*Primary amplification reaction*

9µl of pre-amplification mix (Q5® Hot Start High-Fidelity 2X Master Mix (New England Biolabs, forward cDNA amplifying primer 0.33µM (JP1746, Table S7), reverse cDNA amplifying primer 0.33µM (JP1747, Table S7), primary targeted primers 0.08µM/each (Table S7)) was added to each well and preamplification was carried out with the following cycling protocol: 98ºC/2 min, 20 cycles of 98ºC/10 sec, 60ºC/30 sec, 72ºC 4 min, followed by a final extension at 72ºC/5 min and reactions cooled to 12ºC prior to storage at -20ºC.

*Nested primer reaction*

Following primary amplification, PCR-products were diluted 1:1 with water and mixed. Aliquots of 2µL were transferred to new PCR-plates which already contained 2uL water and reagents for nested targeted PCR were then added using the Mantis system. 8µL of nesting reagents (Q5® Hot Start High-Fidelity 2X Master Mix (New England Biolabs) diluted to 1.5X, nesting primer mix (0.3µM each, Table S7)) were added to each well and cycled as follows; 98ºC/2 min, 20 cycles of 98ºC/10 sec, 65ºC/30 sec, 72ºC/1 min followed by a final extension at 72ºC/5 min and reactions cooled to 12ºC prior to storage at -20ºC.

*Well barcoding*

2µL was transferred from each nested PCR well to a fresh plate containing 4µL of well-specific barcoding primers (each at 1.5 µM; Table S7). 6µL of Q5® Hot Start High-Fidelity 2X Master Mix was added to each well and cycled as follows; 98ºC/2 min, 35 cycles of 98ºC/10 sec, rapid cooling to 80ºC, ramping down to 72ºC at 0.2C/sec, ramping up to 75ºC at 0.2ºC/sec,

75°C/30sec followed by a final extension at 72°C/5 min and reactions cooled to 12°C and kept at 4°C prior to library clean up

*Library clean up, QC, and sequencing*

Barcoded plates were covered with a new plastic lid, edge-sealed with Parafilm, then inverted and centrifuged (200g/room temperature/2 min) to recover and pool barcoded 384-well PCR products. Pools from each plate were cleaned up using a Qiagen PCR clean up kit followed by two rounds of size selection using AMPure beads (Beckman Coulter) at a bead to sample ratio of 0.7:1 (beads to sample). Ligation of barcoded i5 and i7 adapters and sequencing on an Illumina NovaSeq 6000 with an SP flow cell and 2x250bp paired end reads or on an Illumina MiSeq with a v3 3x300bp kit was performed at the Ramaciotti Centre for Genomics – UNSW Sydney. The fastq file names corresponding to each plate are in Table S8.

*Amplicon rescue*

Despite primer optimisation, we observed low read counts for CUX1_p.Q1276* in H198302. This amplicon was rescued with single-plex PCR reactions that repeated primary and nested reactions (Table S7).

For single-plex rescue primary reactions, 2 µL of the original primary amplification product was transferred to fresh 384 well plates containing 8 µL of PCR master mix (0.5U Hot Start Taq DNA Polymerase, 10X Standard *Taq* Reaction Buffer, 2mM $MgCl_2$, 0.3mM dNTP mix (New England Biolabs), and 0.2µM each primary amplification primer) and cycled as follows; 95°C/3 min, 35 cycles of 95°C/20 sec, 60°C/30 sec, 68°C/1 min, followed by a final extension at 68ºC/5 min. Unlike the initial library construction, betaine (Sigma-Aldrich) was included at a final concentration of 1M.

For nested single-plex reactions, 2 µL of amplified product was transferred to fresh 384 well plates containing 8µL of master mix (0.5U Hot Start Taq DNA Polymerase, 10X Standard *Taq* Reaction Buffer, 2mM $MgCl_2$, 0.3mM dNTP mix (New England Biolabs), 0.2µM nesting primer) and cycled as follows; 95ºC/3 min, 35 cycles of 95ºC/20 sec, 65ºC/30 sec, 68ºC/1 min, followed by a final extension at 68ºC/5 min.

Subsequent well barcoding and pooling into libraries was performed as for the initial multiplexed reactions.

**Bioinformatic analyses**

*Code availability*

Code used for bioinformatic analysis can be found at https://github.com/julie-thoms/MDS_amplicons .

*Well demultiplexing, read alignments, and read assignment as wildtype (WT) or mutant (MT)*

Read pairs were demultiplexed into individual wells using the designed sequence barcodes (Table S7). To identify the amplicon, the first 40 bases of read 1 were aligned against all amplicons using seqanpy (https://github.com/iosonofabio/seqanpy). To assign each pair to a wildtype (WT) or mutant (MT) allele, 100 bases of reads (or 60 bp for H198302 CUX1 and H198304 TET2[b]) were aligned to both WT and MT expected sequences and their alignment scores compared. In case of ties, the same was done for read 2. If still tied, the pair was discarded. For each patient, cell type, well, amplicon, and allele, the number of assigned pairs was then computed by simple summation.

*Indexing and assignment of cell types*

Functions for this analysis can be found in the python module index_flow.py. FCS files containing indexed data for each well were imported into python and the embedded compensation matrix applied to the data. For PB-derived cells, populations were well separated for all markers and gating was essentially the same as used for the initial sort. For BM-derived cells, gates were set for each fluorophore based on FMO controls and population shape, except for CD38 which as expected did not resolve into discrete populations, but rather, had cells across a continuum from low to high CD38 expression (Figure S1C). We called the lowest ~10% expressing cells as stem cells ($CD38^{lo}$), and remaining cells as progenitors ($CD38^{+}$). Stem and progenitor cells were then assigned as HSC/MPP ($LIN^{-}$, $CD34^{+}$, $CD38^{lo}$, $CD45RA^{-}$, $CD123^{-}$, $IL1RAP^{-}$**)**, MDS-SC ($LIN^{-}$, $CD34^{+}$, $CD38^{lo}$, [$CD45RA^{+}$ or $CD123^{+}$ or $IL1RAP^{+}$], CMP ($LIN^{-}$, $CD34^{+}$, $CD38^{+}$, $CD45RA^{-}$, $CD123^{+}$), GMP ($LIN^{-}$, $CD34^{+}$, $CD38^{+}$, $CD45RA^{+}$, $CD123^{+}$), or MEP ($LIN^{-}$, $CD34^{+}$, $CD38^{+}$, $CD45RA^{-}$, $CD123^{-}$) (Figure S1C). The assigned cell type for each well in each plate was merged with WT and MT read counts for each amplicon in the corresponding cell.

*Single cell VAFs (scVAFs)*

Functions for this analysis can be found in the module index_haps.py. Single cell VAFS were determined individually for each amplicon. Cells with at least 10 reads for a given amplicon were used for the analysis, and scVAF for each cell type was calculated as the average frequency of the mutant allele across all cells that met the 10 read cut off. Standard error of the mean (SEM) was used to estimate variation in scVAFs across the population of cells.

*Calling haplotypes*

Functions for this analysis can be found in the module index_haps.py. Based on bulk VAFS, we assumed that all the tracked alleles occurred on a single chromosome only (i.e./ individual cells were at most heterozygous for each mutation), and for simplicity called each cell as either WT (mutant allele not detected) or mutated (mutant allele detected). Read count distributions were plotted for each patient, amplicon, and sorted cell type (Figure S4). For each patient, we determined a minimum read count for each amplicon, and individual cells were only included in the haplotype analysis if the read count threshold was reached for all amplicons within that cell. Read count thresholds were selected based on the number of cells available for analysis at each read cut off, with an absolute minimum set at 10 reads for all amplicons. Selected cells were considered mutant for a given allele if the proportion of mutant reads exceeded a predetermined cutoff, which in all cases required the presence of at least 2 mutant reads to call the cell as mutated. We set this minimum to allow for technical limitations such as well carryover during library construction prior to well barcoding. Final read count thresholds and mutant proportion cutoffs used were: H198302 – 10 reads/0.2 mutant proportion cutoff, H198303 – 20 reads/0.1 mutant proportion cutoff, H198304 – 20 reads/0.1 mutant proportion cutoff.

*Plotting*

Plots were drawn in python (v3.8.3) using matplotlib (v3.2.2) and seaborn (v0.10.1). Plotting functions can be found in the modules index_flow.py and index_haps.py and within individual jupyter notebooks.

#### Data availability

Raw sequencing data is available at SRA, accession: PRJNA798507. Flow cytometry data is available at https://flowrepository.org/id/FR-FCM-Z4PR. Full capture sequencing data is available on request from the authors.

#### SUPPLEMENTARY FIGURES

Figure S1 – Gating strategy for isolating individual cell types and assigning cell type based on index sorting data

Figure S2 – Single cell amplicon sequencing library construction

Figure S3 – Characterisation of VAFs in circulating lymphocytes from patient H198304

Figure S4 - Single cell amplicon read distributions

Figure S5 – Bulk and single cell VAFS are highly correlated

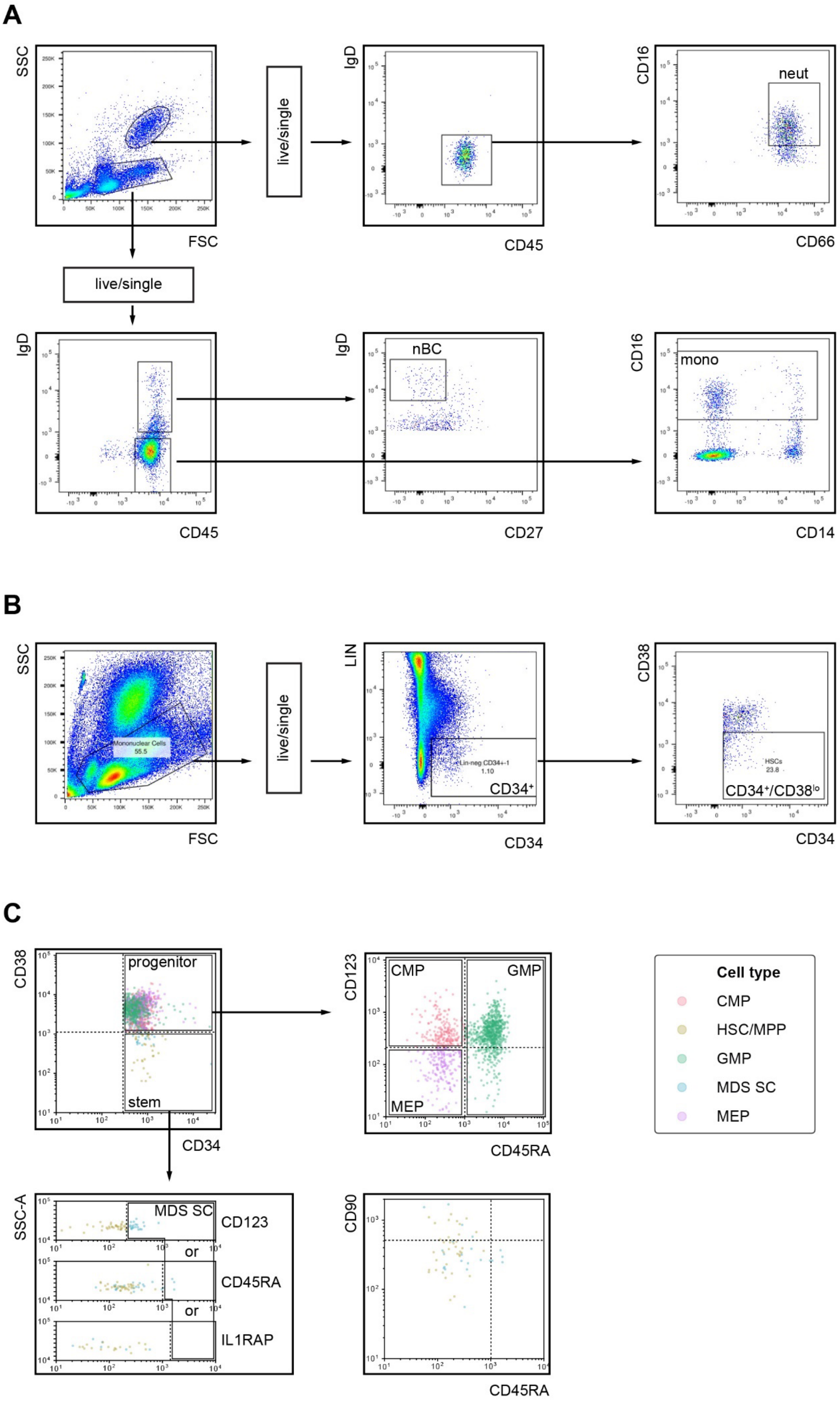
A
SSC
FSC
live/single
IgD
CD45
CD16
neut
CD66
live/single
IgD
CD45
IgD
nBC
CD27
CD16
mono
CD14
B
SSC
Mononuclear Cells
55.5
FSC
live/single
LIN
Lin-neg CD34+-1
1.10
CD34⁺
CD34
CD38
HSCs
23.8
CD34⁺/CD38^lo
CD34
C
CD38
progenitor
stem
CD34
CD123
CMP
GMP
MEP
CD45RA
Cell type
CMP
HSC/MPP
GMP
MDS SC
MEP
SSC-A
MDS SC
CD123
or
CD45RA
or
IL1RAP
CD90
CD45RA

**Figure S1 – Gating strategy for isolating individual cell types and assigning cell type based on index sorting data**

Panels show representative data from patient H198304. **(A)** Sorting strategy for peripheral blood cells. Cells were gated on forward (FSC) and side (SSC) scatter, selected for live (i.e. Zombie$^{lo}$) single cells, then gated as indicated. **(B)** Sorting strategy for bone marrow cells. Cells were gated on FSC and SSC, selected for live single cells, then as lineage negative (LIN$^{-}$) and CD34$^{+}$. For each patient we sorted two to three plates of LIN$^{-}$ CD34$^{+}$ cells, and up to one additional plate where cells were sorted for LIN$^{-}$ CD34$^{+}$ CD38$^{lo}$ to enrich for stem cells. Individual stem and progenitor cell types were resolved using the indexing data for each collected cell. **(C)** Post-hoc BM cell type assignment based on indexing data. Sorted LIN$^{-}$ CD34$^{+}$ cells were computationally called as positive or negative for each surface marker and assigned to cell types based on the combination of markers expressed. For progenitor cells (LIN$^{-}$ CD34$^{+}$ CD38$^{+}$) CD123 and CD45RA were used to resolve CMPs, MEPs, and GMPs. For stem cells (LIN$^{-}$ CD34$^{+}$ CD38$^{lo}$), cells were classed as MDS stem cells (MDS SC) if they were positive for any of the markers CD45RA, CD123 or IL1RAP[6]. Stem cells which did not express any of CD45RA, CD123 or IL1RAP were called as healthy HSC/MPPs. CD123, CD45RA, and IL1RAP distributions within LIN$^{-}$ CD34$^{+}$ CD38$^{lo}$ cells are shown separately. Although the indexing panel included CD90 and was able to distinguish between HSCs and MPPs, numbers of healthy stem cells were generally low and we chose to pool these populations for analysis. *Bottom right:* A representative plot showing CD90 and CD45RA signal for LIN$^{-}$ CD34$^{+}$ CD38$^{lo}$ cells indicates distribution of HSCs (CD90$^{+}$) and MPPs (CD90$^{-}$). Plots show all cells sorted including plates that were not subsequently used for sequencing.

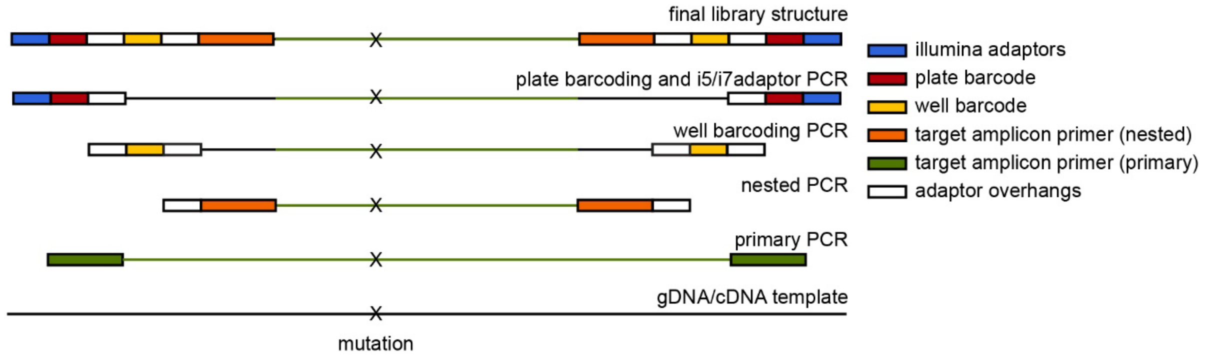


**Figure S2 – Single cell amplicon sequencing library construction**

Schematic showing primary, nested and well-barcoding and plate-barcoding PCR amplification strategies for single cell amplicon libraries. Primary, nested, and well barcoding PCRs were performed on single cells in individual wells, then pools from each 384-well plate barcoded for Illumina sequencing.

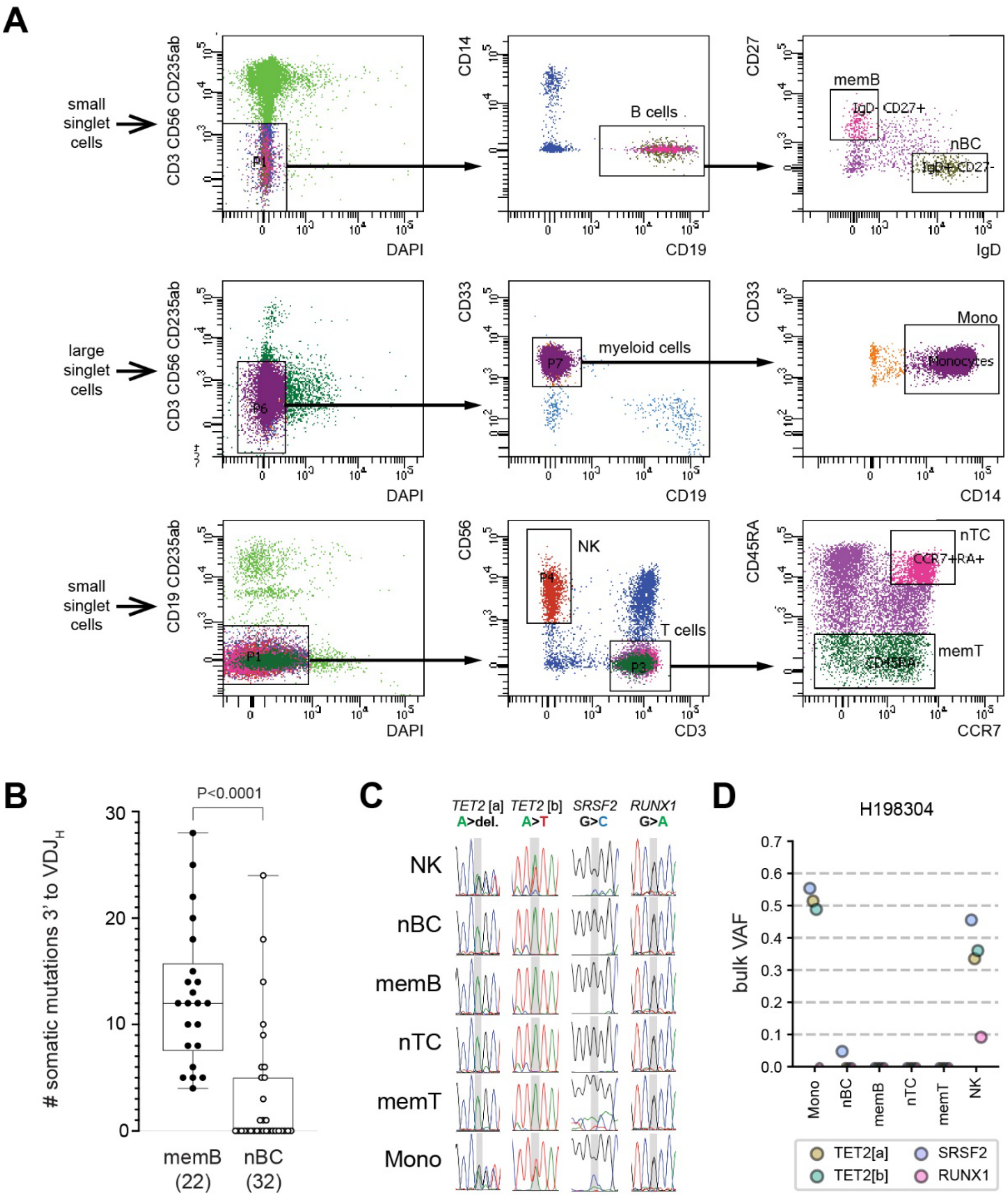
A
small singlet cells
large singlet cells
CD3 CD56 CD235ab
CD19 CD235ab
DAPI
P1
P6
CD14
CD19
B cells
CD33
myeloid cells
P7
CD56
CD3
NK
T cells
CD27
IgD
memB
IgD- CD27+
nBC
IgD+ CD27-
CD14
Mono
Monocytes
CD45RA
CCR7
nTC
CCR7+RA+
memT
B
P<0.0001
### somatic mutations 3' to VDJ
memB (22)
nBC (32)
C
TET2 [a] A>del.
TET2 [b] A>T
SRSF2 G>C
RUNX1 G>A
NK
nBC
memB
nTC
memT
Mono
D
H198304
bulk VAF
Mono
nBC
memB
nTC
memT
NK
TET2[a]
TET2[b]
SRSF2
RUNX1

**Figure S3 – Characterisation of VAFs in circulating lymphocytes from patient H198304**

**(A)** Populations enriched for live (DAPI$^{lo}$) naïve B cells (nBC; CD19+ IgD+ CD27-), memory B cells (memB; CD19+ IgD- CD27+), naïve T cells (nTC; CD3+ CCR7+ CD45RA+), memory T cells (memT; CD3+ CD45RA-), NK cells (CD56+) or monocytes (Mono; CD33+ CD14+) were sorted from an RBC-depleted sample of PB from patient H198304. **(B)** *IgH* VDJ-rearrangements were amplified from the DNA of sorted B cells as described[3], cloned and Sanger sequenced. The frequency of activation-induced *IgH* somatic mutations (insertion, deletion or point mutation) detected within 286bp immediately 3' to unique $VDJ_{H4}$- or $VDJ_{H3}$-rearrangements from $IgD^-$ $CD27^+$ (memB) or $IgD^+$ $CD27^-$ (nBC) B cells. Rearrangements sampled from $IgD^-$ $CD27^+$ B cells universally bore multiple *Ig* somatic mutations consistent with a memory phenotype. 20 of 32 rearrangements sampled from $IgD^+$ $CD27^-$ cells carried zero *IgH* somatic mutations, consistent with a population enriched for naïve B cells, nonetheless minority-contaminated with previously activated (possibly B1) B cells. *p*-value from two-tailed Mann-Whitney test. (**C**) Variant *TET2, SRSF2* and *RUNX1* allele detection by fluorescent Sanger sequencing of bulk cell PCR amplicons. **(D)** Bulk VAFs from the same isolated cell populations.

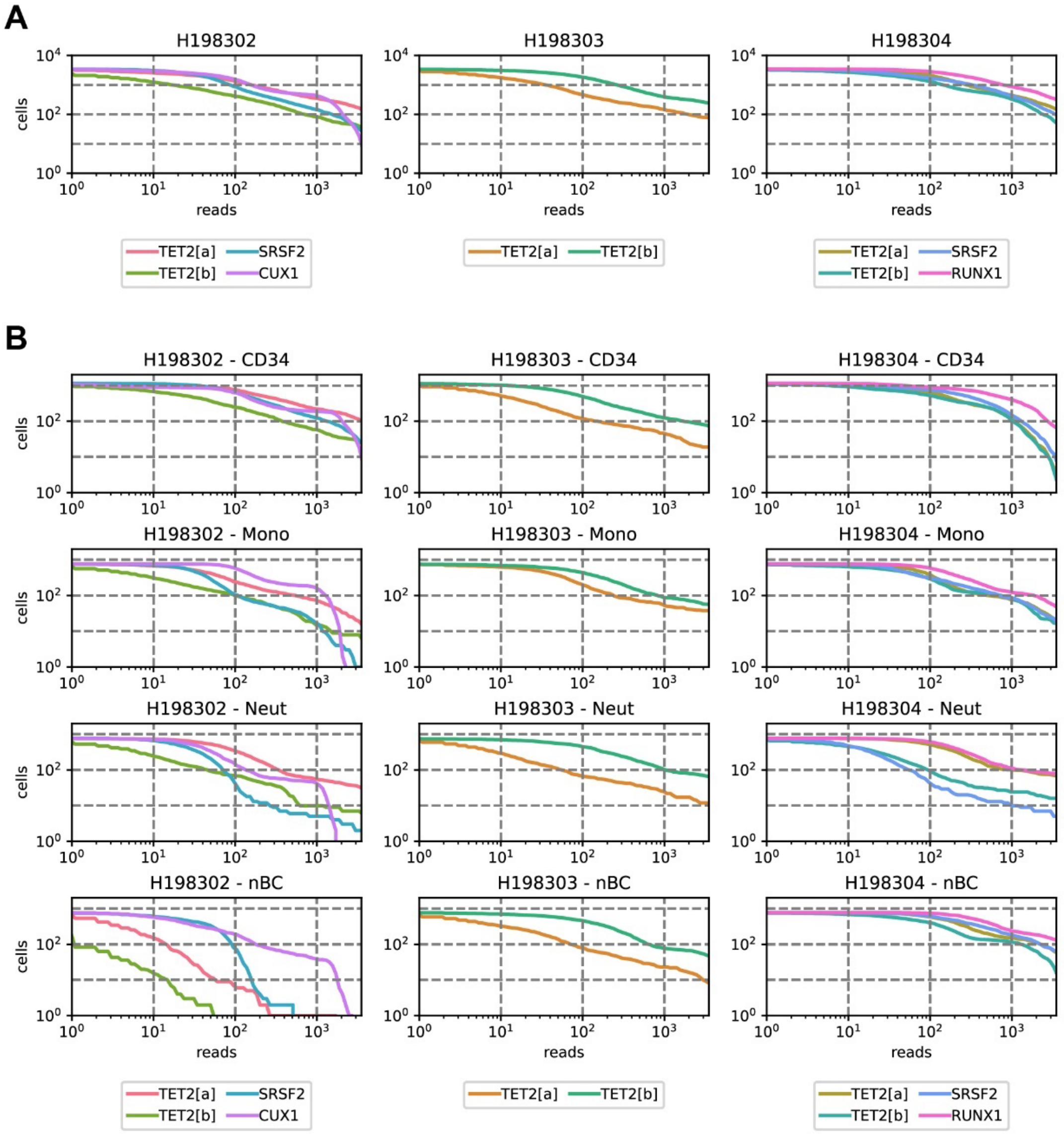


**Figure S4 – Single cell amplicon read distributions**

(A) Plots showing distribution of reads per cell for each amplicon across all cell types in each patient (*L-R:* H198304, H198302, H198303). (B) Plots showing distribution of reads per cell for each amplicon and sorted cell type in each patient (*L-R:* H198304, H198302, H198303).

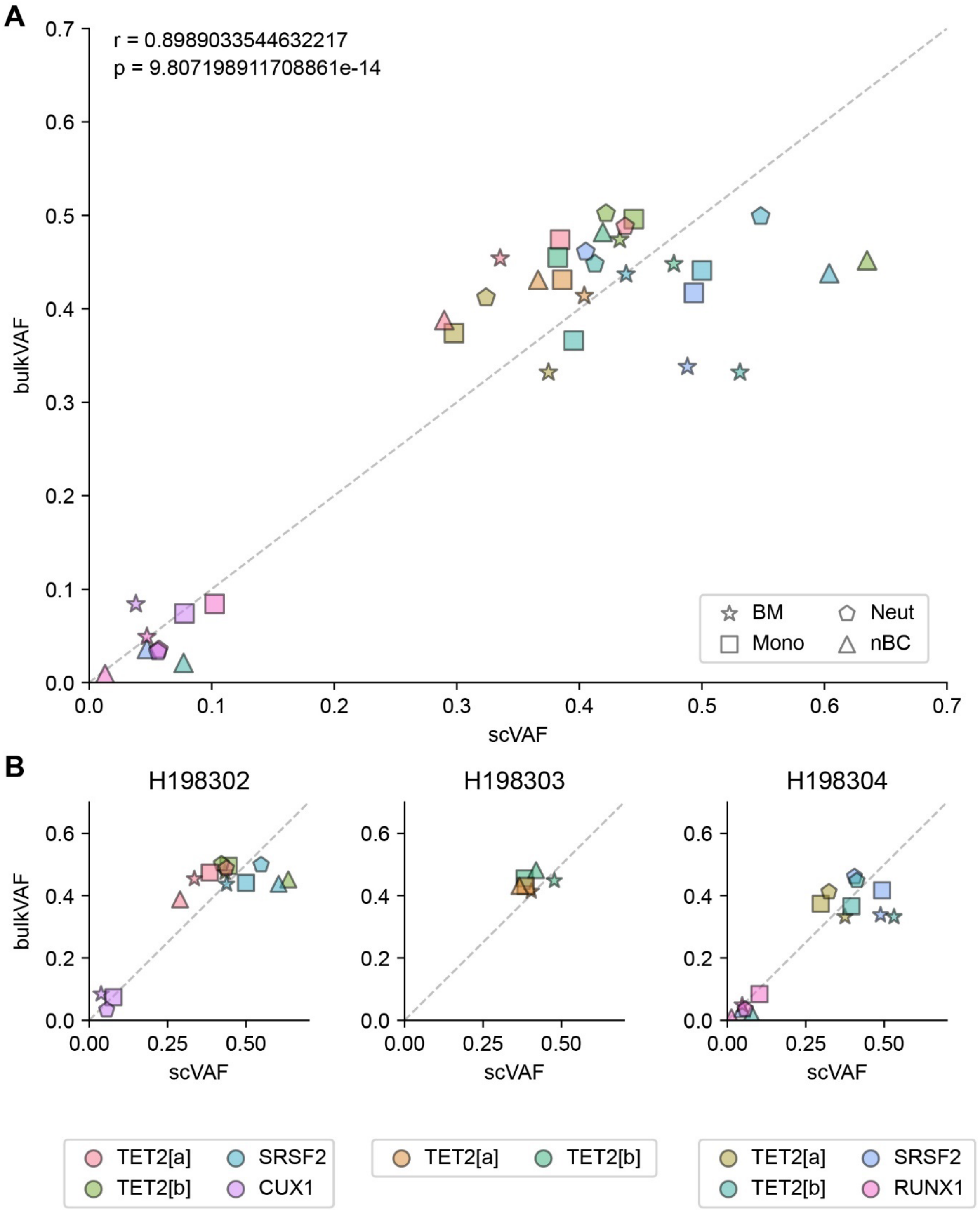

A
r = 0.8989033544632217
p = 9.807198911708861e-14
bulkVAF
scVAF
BM
Neut
Mono
nBC
B
H198302
H198303
H198304
TET2[a]
SRSF2
TET2[b]
CUX1
RUNX1

**Figure S5 – Bulk and single cell VAFS are highly correlated**

**(A)** Composite plot showing all single cell VAFs (scVAFs; x-axis) and bulk VAFs (y-axis) for all patients. VAFs are highly correlated between the two analysis methods (Pearson correlation), dotted line indicates exact concordance. Bulk VAFs were determined in total BM MNCs, while corresponding scVAFs were determined in sorted $CD34^+$ cells. Star – BM/$CD34^+$, square – monocytes, pentagon – neutrophils, triangle – naïve B cells. **(B)** Plots comparing scVAFs and bulk VAFs in each patient (L-R: H198304, H198302, H198303), dotted line indicates exact concordance.
